# *De novo* whole genome assembly of the globally invasive green shore crab *Carcinus maenas* (Linnaeus, 1758) via long-read Oxford Nanopore MinION sequencing

**DOI:** 10.1101/2025.05.19.654951

**Authors:** Jolanda K. Brons, Thomas Hackl, Riccardo Iacovelli, Kristina Haslinger, Sébastian Lequime, Sancia E.T. van der Meij

**Author notes:** these authors contributed equally to this work (Jolanda K. Brons, Thomas Hackl).

## Abstract

Invasive species are rapidly reshaping aquatic ecosystems worldwide at an accelerating pace, with profound ecological and economic impacts. Many crustacean species have demonstrated invasive potential or are already well-established invaders. The green shore crab, *Carcinus maenas*, native to Europe and North Africa, is one of the most successful global marine invaders and is now present on six continents. Although the role of genomics in invasion science is increasingly recognized, genomic resources for brachyuran crabs remain limited, including the notable absence of a reference genome for *C. maenas*. Here we report on a *de novo* whole genome assembly of *C. maenas* via long-read Oxford Nanopore Technology sequencing. The assembly spans 1.09 Gbp across 21,887 scaffolds (N50 = 15 Mbp) with a BUSCO completeness of 98.4%, providing a high-quality resource for future genomic analyses. Additionally, we provide a detailed protocol for obtaining high-quality DNA to successfully sequence brachyuran crabs using a long-read approach, including strategies to address nanopore blockage issues. This new resource expands available genomic data for the species-rich infraorder Brachyura, and provides a valuable foundation for understanding the genetic factors underlying the global invasion success of *C. maenas*, supporting future research in marine invasion genomics.

## Introduction

Invasive species are transforming marine habitats worldwide (Molnar et al., 2008), and the rate at which species occur outside their native ranges has increased considerably over the past few centuries. This increase does not show any sign of saturation, indicating that current efforts to mitigate invasions are inadequate to keep up with the accelerating impacts of globalisation (Seebens et al., 2017). While the drivers of biological invasion are increasingly global in nature, the impacts of invasions are mainly observed at local scales (Molnar et al., 2008). Some non-native species might provide benefits to an ecosystem (Schlaepfer, 2018), while others can cause severe ecological or economic damage, making them invasive (Simberloff et al., 2013). The negative impacts of invasive species range from habitat destruction and disease transmission to displacement or even extinction of native species (Molnar et al., 2008).

The list of the 100 most notorious globally invasive species in the Global Invasive Species Database, compiled by Lowe et al. (2000), includes nine aquatic invertebrates, of which whole genome data is currently available for six. These include three molluscs: the golden apple snail (*Pomacea canaliculata* (Lamarck, 1822)) (Lu et al., 2024), the Mediterranean mussel (*Mytilus galloprovincialis* Lamarck, 1819) (Han et al., 2024), and the zebra mussel (*Dreissena polymorpha* (Pallas, 1771)) (McCartney et al., 2022); an echinoderm, the Northern Pacific seastar (*Asterias amurensis* Lutken, 1871) (Wang et al., 2023); a ctenophore, *Mnemiopsis leidyi* A. Agassiz, 1865 (Ryan et al., 2013); and a crustacean, the Chinese mitten crab (*Eriocheir sinensis*) H. Milne Edwards, 1853 (Cui et al., 2021). Whole genome data is missing for three species: the fishhook waterflea (*Cercopagis pengoi* (Ostroumov, 1891)), the Asian clam (*Potamocorbula amurensis* (Schrenck, 1861)), and the green shore crab (*Carcinus maenas* (Linneaus, 1758)).

These invasive species exhibit traits commonly associated with successful invaders, including broad environmental tolerance, high reproductive output (r-strategists), phenotypic plasticity, effective dispersal, and competitive advantages (Davidson et al., 2011; Weir & Jalice, 2011). Additionally, invasive species may gain additional advantage by escaping natural enemies, although this effect should not be overstated (Colautti et al., 2004). However, the success of many invasive species may rely more on their capacity to adapt through natural selection than on general physiological tolerance or plasticity alone (Lee, 2002). Whole-genome data for invaders are invaluable for advancing invasion genomics (Jaspers et al., 2021) and offer powerful tools to enhance our understanding of biological invasions in aquatic systems (Tepolt, 2015; Bourne et al., 2018; North et al., 2021). For example, comparative whole-genome sequencing in crustaceans demonstrated that many expanded gene families are involved in the environmental tolerance of the red swamp crayfish *Procambarus clarkii* (Girard, 1852) (Xu et al., 2021), whereas the invasion potential of the Chinese mitten crab *E. sinensis* is largely attributed to its strong osmoregulatory capacity and high fertility (Cui et al., 2021). Despite their potential, these resources so far remain underexploited in efforts to predict invasive potential (Kołodziejczyk et al., 2025).

The green shore crab, *Carcinus maenas*, native to Europe and northwest Africa, is one of the world’s most widespread invasive marine species, now recorded from six continents (Carlton & Cohen, 2003; GBIF, 2025; Supplementary Figure 1). Despite its global distribution and ecological impact, no reference genome is currently available for this species. As a generalist predator with a broad diet, *C. maenas* poses serious threats to native biodiversity across its introduced range (Ens et al., 2021). Due to its adaptability and resilience, *C. maenas* is likely to continue expanding into new ecosystems and may trigger secondary invasion waves in areas it already inhabits (Jeffery et al., 2017; Frederich & Lancaster, 2024 - and references therein). The lack of genomic data for *C. maenas* highlights a wider gap in genomic resources for decapod crustaceans (Yuan et al., 2023). Obtaining whole decapod genomes is particularly challenging because they are characterized by their large size and high content of repetitive DNA (Van Quyen et al., 2020; Cui et al., 2021; Rutz et al., 2023). The infraorder Brachyura (true crabs) accounts for almost half of the total diversity in Decapoda, and this group has not only successfully conquered the marine realm, from the deep sea to the intertidal, but is also successful in a multitude of freshwater and terrestrial environments (De Grave et al., 2023). Despite the diversity of this group, with close to 8000 species, and the presence of several notorious invaders and economically important species, only 10 reference genomes across five families are available for brachyuran crabs (NCBI, 2025) with seven more on the way (Wang et al., 2025).

In this study, we present a whole genome assembly of *C. maenas* (family Carcinidae MacLeay, 1838), based on a specimen from within its native range, using long-read sequencing (Oxford Nanopore Technologies (ONT), Oxford, UK). Nanopore technology is a third-generation sequencing method that produces long reads of consistent quality and is recognized as an affordable option for *de novo* sequencing, even for complex genomes (de Lannoy et al., 2017). Although this technology has significant potential to advance genomic research on non-model organisms like aquatic invertebrates, its application is often hindered by challenges in extracting and purifying genomic DNA from many species (Daniels et al., 2023). To address this, we provide a detailed, step-by-step protocol for obtaining sufficient DNA for nanopore sequencing and genome assembly, along with a practical guide to sequencing brachyuran crabs using a MinION, including strategies to overcome pore blockage issues.

## Material and methods

### Specimen collection

Preliminary test runs were performed using an adult male *Carcinus maenas* collected on 26 April 2023 from the marina of Schiermonnikoog, the Netherlands (53.470427, 6.166608). The specimen (Supplementary Figure 2) was preserved in 96% ethanol and stored at –20 °C. Data from these initial runs contributed to the final genome assembly. The whole genome assembly was based on a second adult male *C. maenas*, collected on 11 April 2024 from the harbour of Lauwersoog, the Netherlands (53.410545, 6.208381). This specimen was preserved in DESS solution containing 20% dimethyl sulfoxide (DMSO), 0.25 M ethylenediaminetetraacetic acid (EDTA), and saturated sodium chloride (NaCl), then stored at −20 °C until DNA extraction (Oosting et al., 2020). Both crabs remain vouchered at the University of Groningen.

### Preliminary MinION runs and method optimisation

During initial MinION test runs using muscle tissue from the Schiermonnikoog specimen, consistent pore blockage occurred approximately two hours into sequencing. To address this, we tested various preservation fluids, optimized DNA extraction protocols, and employed both short fragment removal kits and whole genome amplification to eliminate potential contaminants responsible for the blockages. This refined approach, combining improved DNA extraction with two different short fragment removal kits, was subsequently applied to the whole-genome sequencing of *C. maenas*, as detailed below.

The step-by-step protocol for the genomic DNA extraction of *C. maenas* in this study is as follows:

#### 1) Modified Qiagen G/20 Tip DNA Extraction

The Lauwersoog crab specimen was dissected, and approximately 400 mg of muscle tissue from the pereiopods was finely minced. The tissue was suspended in 800 μl of 1x phosphate-buffered saline (PBS, Fisher Scientific, Hampton, USA) and centrifuged at 4000 × g for 1 minute at room temperature. The supernatant was discarded, and the washing step was repeated. The pellet was then resuspended in a solution of 16 μl RNase (Qiagen, Hilden, Germany) and 2 ml G2 buffer (Genomic DNA Buffer Set, Qiagen, Hilden, Germany). Subsequently, 250 μl of proteinase K (>600 mAU/ml, Qiagen, Hilden, Germany) was added, and the mixture was incubated at 50°C for 18 hours.

After incubation, the mixture was centrifuged at 4000 × g for 5 minutes at room temperature. The supernatant was transferred to a new tube, and 1 ml of InhibitEX buffer (Qiagen, Hilden, Germany) was added to remove contaminants (Boughattas et al., 2021). The mixture was incubated at room temperature for 5 minutes and then centrifuged again at 4000 × g for 5 minutes. The resulting supernatant was used for genomic DNA extraction.

Genomic DNA from *C. maenas* was extracted using the ‘Isolation of Genomic DNA from Blood, Cultured Cells, Tissue, Yeast, or Bacteria Using Genomic-tips’ protocol outlined in the Qiagen Genomic DNA Handbook, with the following slight modifications. A Qiagen genomic-tip 20/G (Qiagen Genomic-tip 20/G Kit, Qiagen, Hilden, Germany) was equilibrated with 1.0 ml of QBT buffer (Genomic DNA Buffer Set) and allowed to empty via gravity flow. The *C. maenas* sample was then applied to the prepared genomic-tip 20/G. After the sample passed through, the genomic tip was washed three times with 1.0 ml of QC buffer (Genomic DNA Buffer Set).

The genomic DNA was eluted with 2.0 ml of pre-warmed (50°C) QF buffer (Genomic DNA Buffer Set) and precipitated by adding 0.7 volumes (1.4 ml) of room-temperature (15–25°C) isopropanol (Sigma-Aldrich, Saint Louis, USA). The mixture was gently inverted several times and incubated at room temperature for 5 minutes. It was then centrifuged at 5000 × g for 15 minutes at 4°C. The supernatant was discarded, and the resulting pellet was washed with 1 ml of cold 70% ethanol (4°C) and centrifuged again at 5000 × g for 10 minutes at 4°C. After removing the supernatant, the pellet was air-dried for 10 minutes and resuspended in 60 μl of pre-warmed (50°C) Tris-EDTA (TE) buffer (Sigma-Aldrich, Saint Louis, USA). The resuspension was incubated at 37°C for 1 hour. DNA concentrations were measured using a Qubit Flex Fluorometer (Fisher Scientific, Waltham, USA). The A260/280 and A260/230 ratios were also assessed using a NanoDrop spectrophotometer (Thermo Fisher Scientific, Waltham, USA).

#### 2) Powersoil Pro DNA Extraction

The crab specimen from Lauwersoog was subsampled again, and approximately 400 mg of muscle tissue was finely chopped. The tissue was combined with 800 μl of 1x phosphate-buffered saline (PBS, Fisher Scientific, Hampton, USA) and centrifuged at 4000 × g for 1 minute at room temperature. The supernatant was discarded, and the washing step was repeated twice. DNA extraction was performed using the Powersoil Pro DNA Extraction Kit (Qiagen, Hilden, Germany) according to the manufacturer’s instructions, with the following modifications: 0.25 g of Zirconia beads (Biospec Products, Bartlesville, USA) were added, and the vortexing step (step 2, 10 minutes) was replaced with two 60-second bead-beating cycles at 6.0 m/s using a FastPrep-24™ 5G bead beater (MP Biomedicals, Santa Ana, USA). DNA quality and quantity were assessed as described above.

#### 3) Short Fragment Removal

To enhance the mean read length and investigate pore blockage potentially caused by small reads, a short fragment removal (SFR) step was implemented. This process removed short DNA fragments up to 10 kb (SFR-A) or 25 kb (SFR-B). The SFR-A solution was prepared with 4% PVP 360,000, 1.2 M KCl, and 20 mM Tris-HCl (pH 8), while the SFR-B solution contained 3% PVP 360,000, 1.2 M NaCl, and 20 mM Tris-HCl (pH 8) (Jones et al., 2021). Sixty μl of the *C. maenas* genomic DNA sample was mixed with 60 μl of either SFR-A or SFR-B solution, and the mixture was homogenized by gently tapping the tube. For SFR-B, the mixture was first incubated for 1 hour at 50°C. Both SFR-A and SFR-B samples were then centrifuged at 10,000 × g for 30 minutes at room temperature. The supernatant was carefully removed, and 200 μl of cold 70% ethanol was added, followed by centrifugation at 10,000 × g for 2 minutes at room temperature. This ethanol-washing step was repeated three times. The pellet was subsequently dried for 10 minutes at 50°C and resuspended in 60 μl of pre-warmed (37°C) TE buffer for 20 minutes at 37°C. DNA quality and quantity were assessed as described above.

#### 4) Whole Genome Amplification

To address persistent pore blockage in the MinION flow cell during sequencing, several approaches were tested to remove potential contaminants, including alternative DNA extraction methods, incorporation of an InhibitEX step, and short fragment removal (as previously described). Despite these efforts, the blockage issue persisted. Whole genome amplification (WGA) was therefore performed on the *C. maenas* sample (modified G/20 tip DNA extraction + SFR-A kit) using the REPLI-g Mini Kit (Qiagen, Hilden, Germany), following the manufacturer’s instructions, in an effort to eliminate the source of the blockage. DNA concentrations and purity ratios were again assessed using a Qubit Flex Fluorometer and a NanoDrop spectrophotometer.

#### 5) Library Preparation and MinION Sequencing

Genomic DNA libraries for *C. maenas* were prepared using the SQL-LSK114 Ligation Sequencing Kit V14 (ONT). Library preparation started with 1 μg of *C. maenas* DNA, quantified using a Qubit Flex Fluorometer. Following the ‘Ligation Sequencing gDNA V14’ protocol provided by ONT, the process included a DNA repair and end-prep, adapter ligation, and clean-up step. DNA concentrations were measured after each step using the Qubit Flex Fluorometer. The sequencing flow cell (R10.4.1, FLO-MIN114, ONT) was primed and loaded with 20–60 ng of *C. maenas* DNA per run (Supplementary Table 1). Sequencing was performed on a MinION device (ONT), basecalling and demultiplexing were performed with MinKNOW/Dorado at “super-accurate basecalling, 400 bps”.

#### 6) Pore Blockage and Sequencing Optimization

Both the modified G/20 tip and Powersoil Pro DNA extraction methods, combined with either SFR-A or SFR-B or the whole genome amplification sample (modified G/20 tip DNA extraction + SFR-A kit), produced sufficient DNA for sequencing (20–100 ng/μl). The short fragment removal kits (SFR-A and SFR-B) effectively reduced the number of small reads in the samples. However, none of the tested approaches, including whole genome amplification, were able to prevent pore blockage, which consistently occurred approximately two hours into sequencing.

Interestingly, a pore scan revealed that some pores reopened after blockage, and a substantial increase in active pores was observed following the use of the Flow Cell Wash Kit (ONT), which contains DNase. This suggests that pore blockage may not be caused by contaminants, but by structural properties of the DNA itself. To mitigate blockage, we implemented a sequencing protocol that included a pore scan every 30 minutes during the run and a flow cell wash every 2 hours (Supplementary Table 1).

### Genome Assembly and Curation

The read data was assembled with Flye v2.9.2 (Kolmogorov et al., 2019) specifying high-quality nanopore reads as input (`--nano-hq`), and the assembly was polished with medaka v1.11.3 (https://github.com/nanoporetech/medaka).

Due to difficulties in the sequencing process and the resulting limited amount of sequencing data, we generated two initial assemblies. The first assembly used only the data from the Lauwersoog specimen, while the second combined data from this specimen with an additional 10% of reads from the Schiermonnikoog specimen obtained during the test phase. Although combining data from multiple specimens is not generally recommended, since it can introduce small sequence and structural variations that may negatively impact assembly quality, comparison of the statistics of the two assemblies indicated that the combined assembly had better contiguity and completeness. Based on this, we chose to proceed with the combined assembly for all subsequent analyses.

Because a complete mitochondrial genome was not detectable in our draft assembly, we used a targeted approach to assemble it. Mitochondria of crabs have a lower GC content than the nuclear genome, and mitochondrial DNA is present in high abundance in muscle tissue. We therefore extracted reads from our data with a GC content below 35% and a minimum length of 10 kbp and generated an assembly from 10% of the resulting reads. Using mitochondrial-encoded cytochrome c oxidase subunit I (COI) protein sequences obtained from Interpro (IPR000883) as a marker (Blum et al., 2025), we identified a mitochondrial candidate contig using diamond blastx v2.19 (Buchfink et al., 2023) in the target assembly. Whole-genome self-alignment of the candidate sequence revealed that its palindromic structure was an artifact of the assembly process. We pruned the duplicated region to obtain the final mitochondrial sequence. Using this curated sequence as a reference, we removed remaining mitochondrial fragments from the whole genome draft based on minimap2 v2.28 (Li, 2018) mappings before adding the curated mitochondrial sequence to the whole genome assembly.

To optimize our assembly in terms of contiguity, we used homology-based reference scaffolding to organize smaller contigs into larger scaffolds. Specifically, we used the homology scaffolder RagTag (Alonge et al., 2022), leveraging the highly contiguous genome assembly of the mud crab *Scylla paramamosain* Estampador, 1950 as reference (RefSeq accession: GCF_035594125.1).

### Assembly Statistics and Quality Assessment

To assess potential contamination of the assembly, we employed two complementary approaches. First, we ran the deep learning classifier DeepMicroClass v1.0.3 (Hou et al., 2024) for a broad classification of sequences into eukaryotes, prokaryotes and viruses. Second, we used MMSeq2 homology-based `easy-taxonomy` workflow (Mirdita et al., 2021) with UniRef50 (The UniProt Consortium, 2025) as database to assign each contig a last common ancestor classification based on sequence homology.

To estimate assembly completeness, we used Compleasm v0.2.6 (Huang & Li, 2023). To identify low-complexity regions within the sequences, we employed tantan v26 (Frith, 2011). Additionally, we gathered basic assembly statistics—such as sequence length and base composition—using SeqKit (Shen et al., 2024).

## Results

After optimizing genomic DNA extraction and implementing the previously described sequencing protocol, we successfully generated 11.1 Gbp of *C. maenas* DNA sequence data using five MinION flow cells, achieving an average read N50 of 6.11 kbp. This dataset was supplemented with an additional 1.2 Gbp of data generated from the initial test sequencing runs on the Schiermonnikoog specimen. The whole genome of *C. maenas* was assembled *de novo* and polished, generating an initial contig assembly of 965 Mbp, comprising 34,569 contigs and an NG50 of 60 kbp. We further scaffolded our draft contigs based on homology to the mud crab (*Scylla paramamosain)* and curated the resulting scaffolds to ensure high quality (see Material and Methods). We obtained a final genome assembly of 1.09 Gbp in size, comprising 21,887 scaffolds and an NG50 of 15 Mbp (i.e. more than 50% of the genome is contained in scaffolds larger than 15 Mbp).

A closer examination of the number and size distribution of the assembled sequences reveals a bimodal pattern (Figure 1a). Approximately 80% of the genome is represented by chromosome-scale scaffolds, while the remainder consists predominantly of shorter sequences ranging from 1 to 100 kbp. This distribution is consistent with our reference-based scaffolding approach, where some contigs were either unplaced or placed ambiguously due to limited or conflicting homology information.

**Figure 1.**
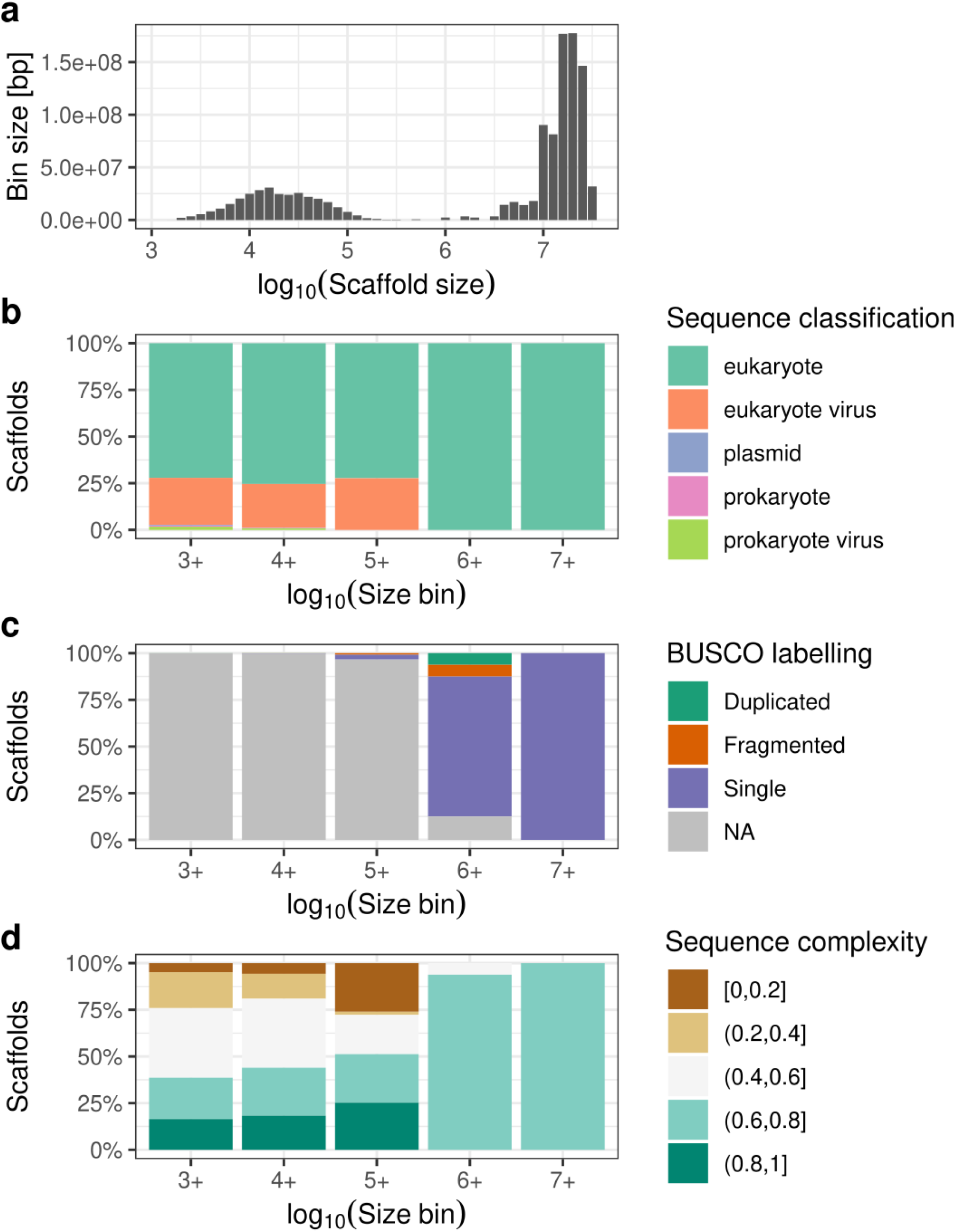
Overview of assembly composition and quality metrics. (**a**) Size distribution histogram of assembled contigs and scaffolds. (**b**) Broad classification profiles across contigs/scaffolds of different length bins (minimum sizes: 1kb, 10kb, 100kb, 1Mb, 10Mb), with colors indicating the predicted source. (**c**) Distribution of universal eukaryotic marker genes (BUSCOs) across the same contigs/scaffolds length bins. (**d**) Sequence complexity profiles across contigs/scaffolds length bins with colors indicating the fraction of sequences identified as low complexity by the tantan algorithm.

Assessment of contamination via two complementary approaches indicated negligible levels of sequences predicted to derive from sources other than the genuine *C. maenas* genome (Figure 1b, Supplementary Table 2 Table “mmseq-report”). Using broad deep-learning-based classification, scaffolds larger than 1 Mbp were exclusively classified as eukaryotic, while ¾ of smaller scaffolds were classified as eukaryotic and ¼ as eukaryotic viruses. In the majority, the latter sequences represent retrotransposons that are expected to be abundant in this genome. Consistently, homology classification placed 89% of sequences with last common ancestors along the Brachyura lineage, with less than 2% of sequences potentially assigned to other kingdoms (1.8% fungi, <1% plants, bacteria and viruses). Completeness estimates based on lineage-specific universal eukaryotic marker genes indicate a highly complete genome assembly, with 98.4% of markers detected (Figure 1c). Most marker genes were assembled at full length and located within large scaffolds. Additionally, the low level of duplicated markers supports the assembly’s high contiguity and suggests minimal structural contamination.

Finally, assessment of the sequence complexity revealed expected high levels of sequences with repetitive nature among the unscaffolded short contigs (Figure 1d). The presence of repetitive regions in these contigs is consistent with both the classification of a significant fraction of these contigs as putative retrotransposons and the apparent difficulty of scaffolding these contigs unambiguously into a longer scaffold context, thereby contributing to the bimodal sequence length distribution observed.

While our initial draft assembly did not contain a detectable mitochondrial genome, we recovered a single mitochondrial contig of 15,474 bp using a targeted assembly approach. The obtained mitochondrial genome matches those of three related crabs (*Scylla paramamosain* - JAHFWG010003966.1; *Callinectes sapidus* Rathbun, 1896 - NC_012572.1, *Portunus trituberculatus* (Miers, 1876) - NC_005037.1) in size and divergence and appears syntenic in its overall genomic organization (Supplementary Figure 3).

## Discussion

Here we present a high-quality genome assembly of the green shore crab, *Carcinus maenas*, with a total size of approximately 1.09 Gbp. The assembly is highly complete, containing 98.4% of universal eukaryotic markers, and achieves high contiguity through homology-based scaffolding. This genome fills an important gap within the Brachyura, bringing the total number of publicly available true crab genomes to eleven.

Despite its ecological significance as an invasive species and its widespread use as a model organism across multiple research fields, particularly ecotoxicology (Leignel et al., 2014; Rodrigues & Pardal, 2014), a complete genome for *C. maenas* was not previously available. Earlier sequencing efforts in 2016 produced a fragmented draft assembly covering only 36% of the estimated genome size, hindered primarily by the genome’s high repetitive content (Verbruggen, 2016). Our assembly thus provides a valuable foundation for future studies on the genomic basis of invasion success and supports the species’ utility as a model organism in other lines of research.

Our *C. maenas* specimens were sequenced using an Oxford Nanopore R10.4.1 platform, which offers high-accuracy long reads suitable for de novo genome assembly (Guiglielmoni & Schiffer, 2024). However, we faced recurring pore blockage approximately every two hours during sequencing. Similar challenges have been reported in other decapods, such as the tiger prawn (*Penaeus (Penaeus) monodon* Fabricius, 1798). The authors attributed these blockages to the formation of complex secondary structures during sequencing and considered them irreversible (Van Quyen et al., 2020). In contrast, we successfully mitigated these issues by applying a Flow Cell Wash Kit every two hours, reopening many blocked pores and extending sequencing runs. Similar strategies were required in sequencing the subtidal gastropod *Kelletia kelletii* (Forbes, 1852), where additional cleanup and DNA purification steps were necessary to overcome similar nanopore sequencing challenges (Daniels et al., 2023).

To facilitate genomic research in brachyuran crabs and other non-model marine invertebrates, we provide a detailed protocol for long-read sequencing using Oxford Nanopore’s MinION platform, from DNA extraction to strategies for managing pore blockage. This resource highlights the importance of methodological optimization in generating high-quality genomic data for ecologically and evolutionarily important marine invertebrate taxa.

## Data availability

Raw sequence data and the assembled, curated genome presented here have been deposited at the European Nucleotide Archive under the project accession PRJEB86588.

## Acknowledgements

We thank Britas Klemens Eriksson, Jorn Claassen, Yichen Liu, Lea Simon (all University of Groningen), Bastian Reijnen and two little girls for providing us with *C. maenas* specimens for (preliminary) genetic analyses and testing of sequencing protocols. We furthermore thank the Center for Information Technology of the University of Groningen for their support and for providing access to the Hábrók high-performance computing cluster.

## Notes

### Competing Interest Statement

The authors have declared no competing interest.

