## Supplementary Figures 1-3 for "*De novo* whole genome assembly of the globally invasive green shore crab *Carcinus maenas* (Linnaeus, 1758) via long-read Oxford Nanopore MinION sequencing"

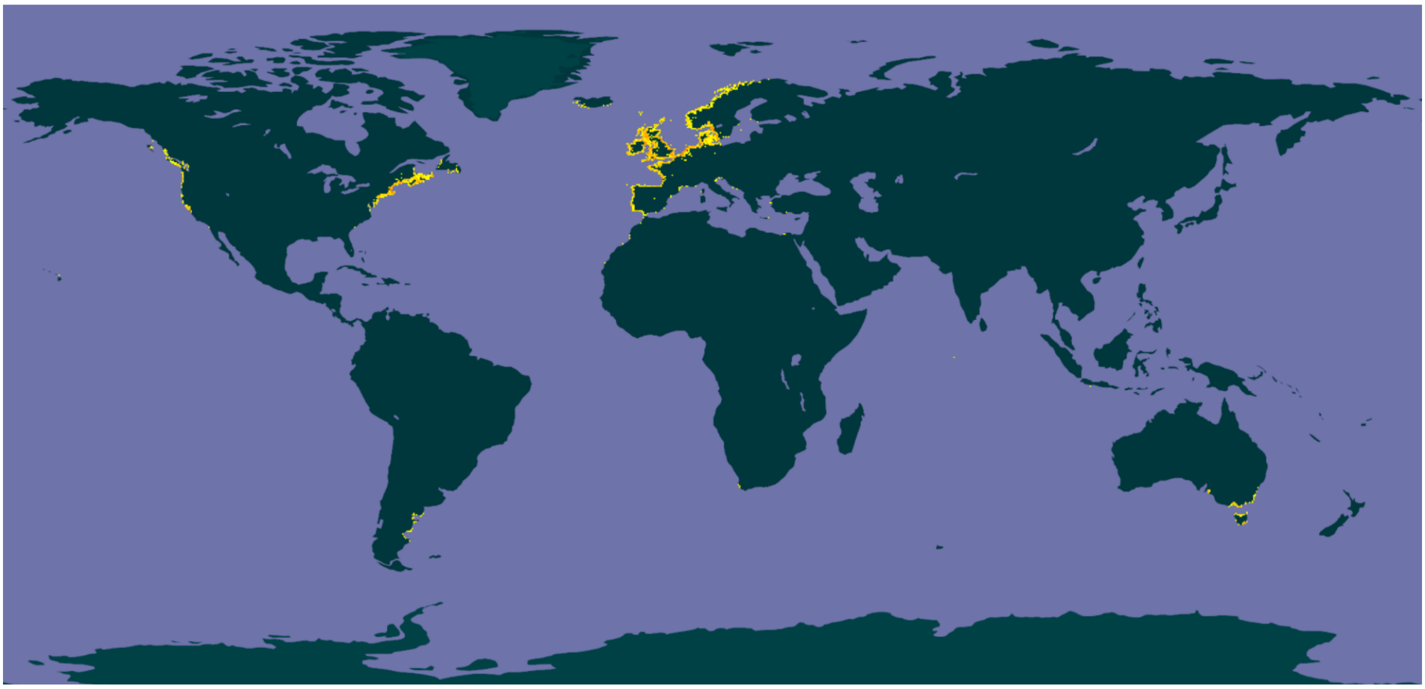

**Supplementary Figure 1.** Occurrence of *C. maenas* (marked as yellow dots) based on GBIF occurrence data [download Dec 2024).

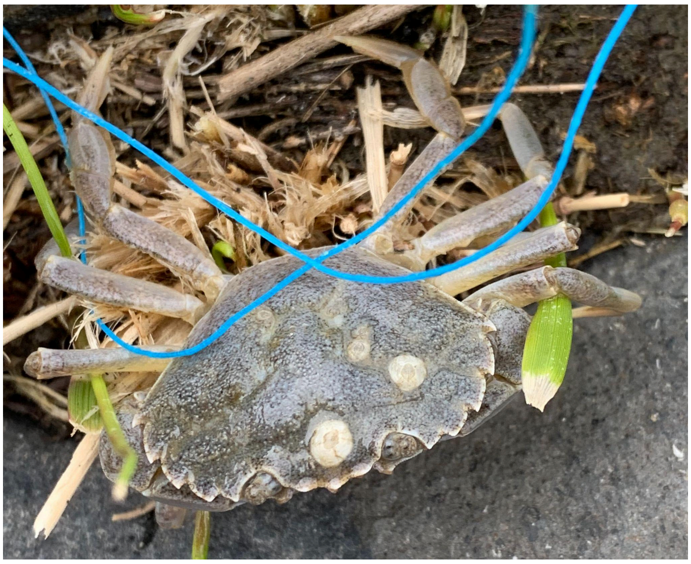

**Supplementary Figure 2.** *Carcinus maenas* specimen collected from the Schiermonnikoog marina using a line with mussel bait.

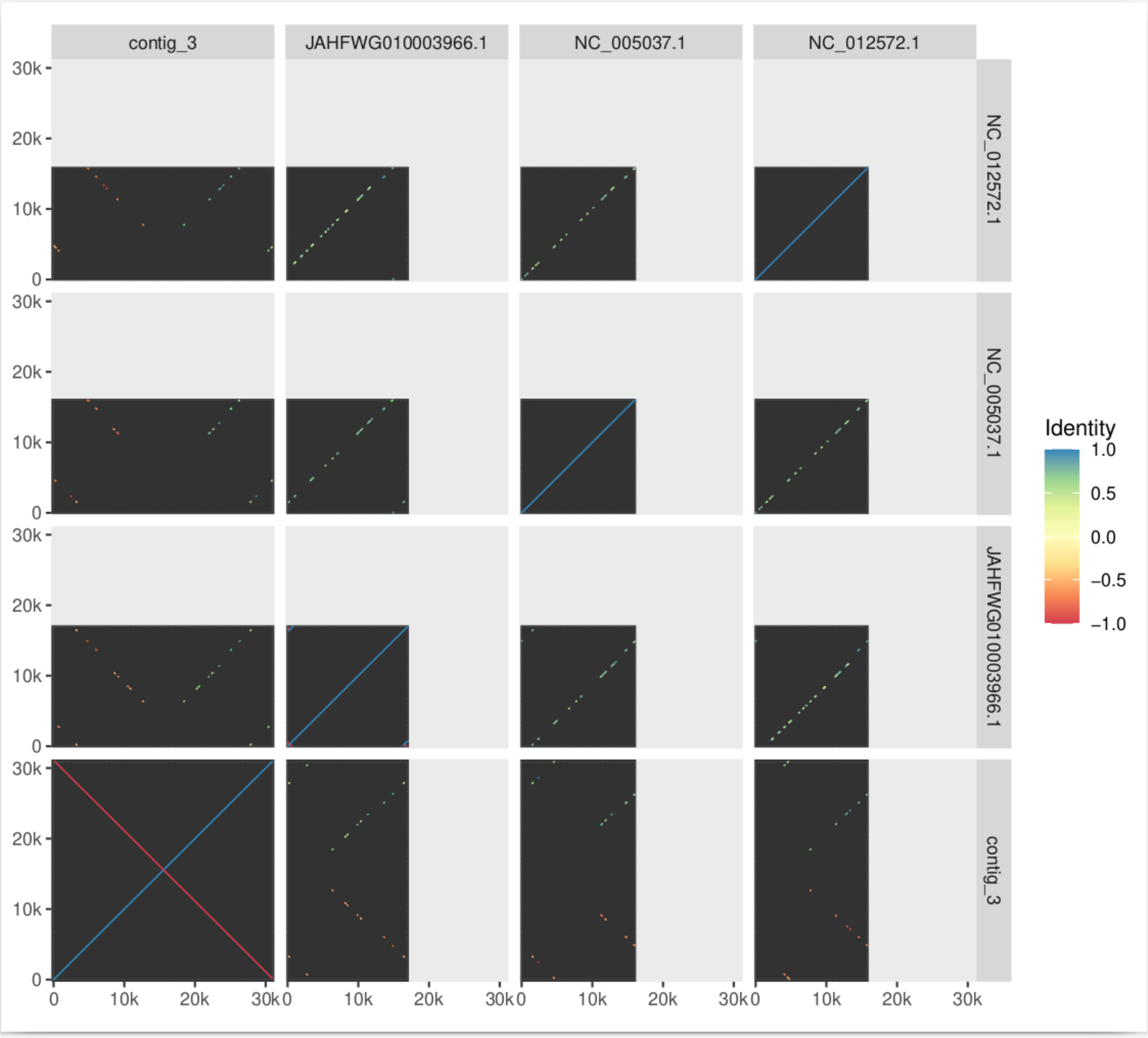


**Supplementary Figure 3. Whole genome alignment of mitochondrial candidate contig with close references.** Visual representation of all-vs-all whole genome alignments of the three mitochondrial genomes of *Scylla paramamosain* (JAHFWG010003966.1), *Callinectes sapidus* (NC_012572.1), and *Portunus trituberculatus* (NC_005037.1) with the mitochondrial genome candidate contig obtained for *C. maenas* (contig_3) in this study. Regions of sequence similarity between two genomes are indicated by colored lines in the plot, with levels of similarity and direction of the alignment indicated by the color scale (blue: same direction, 100% similarity, yellow: low similarity, red: opposite direction, 100% similarity). The plot shows that the *C. maenas* contig consists of two reverse-complement copies of the actual mitochondrial genome sequence – an assembly artifact caused by the terminal internal inverted repeat structure typically found in mitochondria. This assembly artifact was resolved through manual curation as part of the assembly quality control, and a corrected version of the mitochondrial genome was deposited.
